# Benefits of Collisional Cross Section Assisted Precursor Selection (caps-PASEF) for Cross-linking Mass Spectrometry

**DOI:** 10.1101/2020.04.16.044859

**Authors:** Barbara Steigenberger, Henk W.P. van den Toorn, Emiel Bijl, Jean-François Greisch, Oliver Räther, Markus Lubeck, Roland J. Pieters, Albert J.R. Heck, Richard A. Scheltema

**Affiliations:** Biomolecular Mass Spectrometry and Proteomics, Bijvoet Center for Biomolecular Research and Utrecht Institute for Pharmaceutical Sciences, Utrecht University, Padualaan 8, 3584 CH Utrecht, The Netherlands; Netherlands Proteomics Centre, Padualaan 8, 3584 CH Utrecht, The Netherlands; Bruker Daltonik GmbH, Fahrenheitstrasse 4, 28359 Bremen, Germany; Department of Chemical Biology & Drug Discovery, Utrecht Institute for Pharmaceutical Sciences, Utrecht University, 3508 TB Utrecht, The Netherlands

**Keywords:** Crosslinking Mass Spectrometry, timsTOF, XL-MS, PhoX, XlinkX

## Abstract

Ion mobility separates molecules in the gas-phase on their physico-chemical properties, providing information about their size as collisional cross-sections. The timsTOF Pro combines trapped ion mobility with a quadrupole, collision cell and a time-of-flight mass analyzer, to probe ions at high speeds with on-the-fly fragmentation. Here, we show that on this platform ion mobility is beneficial for cross-linking mass spectrometry (XL-MS). Cross-linking reagents covalently link amino acids in close proximity, resulting in peptide pairs after proteolytic digestion. These cross-linked peptides are typically present at low abundance in the background of normal peptides, which can partially be resolved by using enrichable cross-linking reagents. Even with a very efficient enrichable cross-linking reagent, like PhoX, the analysis of cross-linked peptides is still hampered by the co-enrichment of peptides connected to a partially hydrolyzed reagent – termed mono-linked peptides. For experiments aiming to uncover protein-protein interactions these are unwanted byproducts. Here, we demonstrate that gas-phase separation by ion mobility enables the separation of mono-linked peptides from cross-linked peptide pairs. A clear partition between these two classes is observed at a CCS of 500 Å^2^ and a monoisotopic mass of 2 kDa, which can be used for targeted precursor selection. A total of 50 - 70% of the mono-linked peptides are prevented from sequencing, allowing the analysis to focus on sequencing the relevant cross-linked peptide pairs. In applications to both simple proteins and protein mixtures and a complete highly complex lysate this approach provides a substantial increase in detected cross-linked peptides.

---

The folding of proteins, resulting in structural features that enable them to function and form complexes with other proteins, is one of the major driving forces in highly sophisticated cellular behavior. Misfolding and/or gain or loss of interactions to other proteins can lead to major dysfunction and potentially severe diseases(1, 2). Intimate knowledge of the structural details behind protein structures and interactions is of the utmost importance to develop novel treatments to interfere with these dysfunctions. Even though the study of protein structure is dominated by techniques like NMR, crystallography and cryo-EM, structural proteomics techniques driven by mass spectrometry have an increasingly important, integrative role to uncover new details not achievable by the conventional techniques. For example, information on proteoforms (i.e. protein sequences and post-translational modifications) are typically not apparent with a technique like cryo-EM but are readily accessible by structural proteomics(3). At the same time, spatial information within and between proteins can be obtained by the use of cross-linking mass spectrometry (XL-MS)(4–8).

XL-MS typically uses small homo-bi-functional chemical reagents that irreversibly connect amino acids in close structural proximity. Most commonly highly reactive NHS-esters, which primarily capture the sidechains of lysines are used for this purpose. After reduction, alkylation and proteolytic digestion of the cross-linked proteins, three different products are formed: unmodified peptides, peptides with a quenched linker attached termed “mono-link” peptides and the desirable two peptides covalently connected by the cross-linking reagent termed “cross-link” peptides. Cross-linked peptides provide information on protein tertiary structure in the form of intra-links (two peptides from the same protein) and protein quaternary structure in the form of inter-links (two peptides from different proteins). As the reaction efficiency for cross-linking is estimated to be about 1-5%, and relatively few lysine pairs are found to be in sufficiently close proximity to be cross-linked, only 0.1% of the sample actually consists of cross-linked peptides, which substantially hampers their detection(9–11). To focus the analysis, extensive pre-fractionation of the peptide mixture is commonly employed prior to the LC-MS measurement(s), using chromatographic techniques such as strong cation exchange (SCX) or size exclusion chromatography (SEC). However, reagents with an enrichment handle directly attached have emerged capable of removing the high background of normal peptides and uniquely enrich for modified peptide products (mono-linked and cross-linked peptides). For this purpose, conventionally a biotin handle is used – either directly attached to the reagent or introduced after the cross-linking reaction by a click-reaction. One of the downsides of using biotin as enrichment handle is that its high affinity binding to streptavidin prevents efficient elution from the enrichment beads. Recently, we developed and introduced a novel enrichable cross-linking reagent, PhoX, which is decorated with a phosphonic acid moiety directly attached on the cross-linking reagent(9). This moiety is a stable mimic of a phosphate group and can therefore be enriched efficiently by IMAC-based techniques originally developed for phosphorylated peptides. Competing molecules for the affinity enrichment, such as phospho-peptides and nucleic acids, can selectively be removed by using a phosphatase and/or benzonase, as PhoX remains stable under these conditions. With the PhoX enrichment handle, we increased the enrichment efficiency by up to 300x with 97% specificity, leading to excellent cross-link identification. The approach is however not yet focusing solely on the desired cross-linked peptides, as the sample still contains approximately 60% of the less informative mono-linked peptides.

With ion mobility mass spectrometry (IMMS) ions are separated over a time-frame of 10 - 100 ms by their collisional cross section (CCS, Ω) (12, 13), which is based on their size, shape and charge. Ion mobility separation (IMS) devices are typically installed between the liquid chromatography (LC) system and the mass analyzer. It has been demonstrated that ions eluting from an IMS device can efficiently be sampled with TOF analyzers, as these devices have the high acquisition rates – in the range of 10 kHz – required for this fast separation technique. Different conceptions of IMS are currently applied in the field of mass spectrometry, with trapped ion mobility separation (TIMS) featuring several desirable properties, such as small size, low voltage requirements and highly efficient ion utilization. In TIMS, ions are balanced in an electrical field against a constant gas stream allowing ions to be trapped and stored at different positions in the ion tunnel device. After trapping, mobility-separated ions can be released from the TIMS device by lowering the electrical potential and can subsequently be transferred to a mass analyzer. Low mobility ions with large CCS values are eluted first from the TIMS device, followed by high mobility ions with smaller CCS values(14, 15). As cross-linked peptides consist of two peptides connected by the cross-linking reagent, their size and shape typically differs from non-modified and mono-linked peptides and therefore we hypothesized that the TIMS device connected to a TOF analyzer could be an excellent candidate for the required extra level of separation (Fig. 1A).

**Figure 1.**
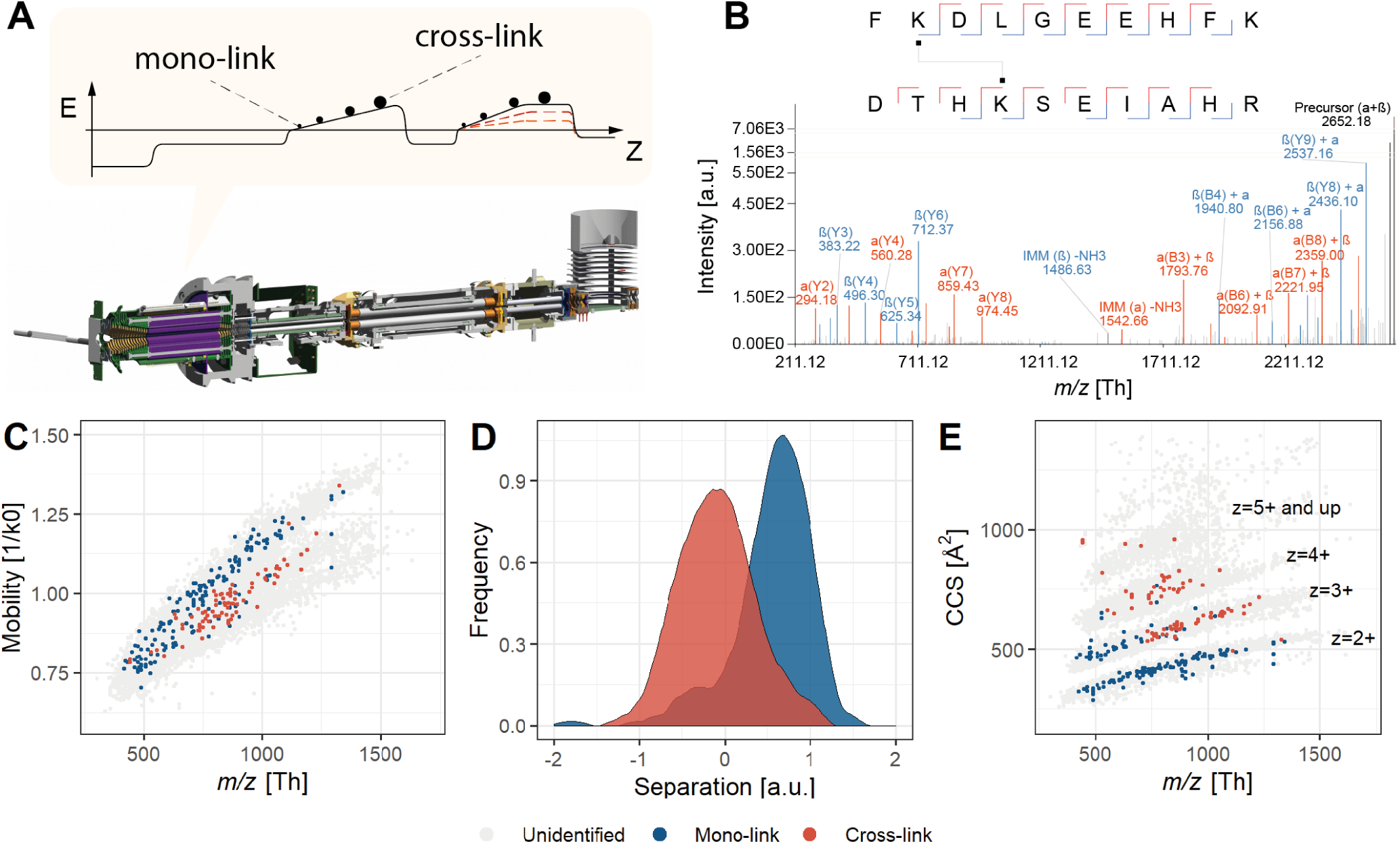
Integration of ion mobility into cross-linking MS. **(A)** Instrument overview with the conceptual operation of the TIMS device separating the PhoX-enriched cross-linked sample into mono-linked and cross-linked peptides. **(B)** Deisotoped tandem mass spectrum demonstrating the fragmentation performance with stepped HCD fragmentation on the timsTOF Pro. **(C)** Distribution of ions signals (originating from PhoX cross-linked BSA) for *m/z* (Th) versus Mobility (1/K0) for all classes of ions (‘Unidentified’: likely noise; ‘Mono-link’: 192; ‘Cross-link’: 80). The legend for the color-coding is provided at the bottom of the figure. **(D)** Physical separation of mono-linked from cross-linked peptides in mobility space. (E) Distribution of *m/z* (Th) versus Collision Cross Section for all classes of ions.

Here, we describe the first application of XL-MS on the timsTOF Pro using the efficiently enrichable cross-linker PhoX(9). We demonstrate, following careful optimization of the parameters, that the system has the sensitivity to detect and identify the typically difficult to interpret cross-linked peptide spectra with its ability to produce high quality fragmentation spectra (Fig. 1B). The TIMS device physically separates the mono-linked and cross-linked peptides, effectively providing the extra dimension of separation required for improved level of detection. Furthermore, we introduce a novel acquisition strategy termed caps-PASEF (Collisional Cross Section Assisted Precursor Selection), which makes use of CCS information to make an easy-to-use a-priori distinction between molecules of interest and demonstrate the performance on standard protein mixture and a complex sample of proteins from a full cellular lysate.

## Materials and Methods

### Cross-linking reagent

A batch of the cross-linking reagent PhoX was synthesized as previously described(9) and freshly dissolved at a concentration of 50 mM in anhydrous DMSO. This solution was divided in separate aliquots and stored in Eppendorf tubes at −20 °C. Each aliquot was for one-time-use only, as the reactive NHS-esters of PhoX can potentially hydrolyze. Prior to opening an aliquot, slow equilibration to room temperature is required to avoid additional water in the solution.

### Synthetic peptides

PhoX was added to synthetic peptides (10 µL, 5 mM in 1xPBS, sequence: Ac-AAAAKAAAAAR-OH) at a final concentration of 2 mM. The cross-linking reaction was incubated for one hour at room temperature and then halted by addition of 5 µL Tris·HCl (100 mM, pH 8). After desalting with Sep-Pak *C*_18_, the peptide mixture was directly infused into the Bruker timsTOF Pro. Ions with masses corresponding to cross-linked or mono-linked peptides were manually isolated and subjected to increasing HCD energy.

### Cross-linking and digestion of proteins

Proteins were incubated with PhoX for 45 min at room temperature (buffer conditions specified below). The cross-linking reaction was quenched by addition of Tris·HCl (100 mM, pH 7.5) to a final concentration of 10 mM. Residual cross-linking reagent was removed by size-cut-off filters (Vivaspin 500K 10kDa MWCO centrifugal filter units) with three volumes of Tris·HCl (100 mM, pH 7.5) or by acetone precipitation. Cross-linked proteins (in 50 mM Tris·HCl, pH 7.5) were reduced with DTT (final concentration of 2 mM) for 30 min at 37 °C, followed by alkylation with IAA (final concentration of 4 mM) for 30 min at 37 °C. This reaction was quenched by addition of DTT (final concentration of 2 mM). Then, the sample was digested by incubating with a combination of LysC (1:75 enzyme to protein) and Trypsin (1:50 enzyme to protein) for 10 h at 37 °C, after which formic acid (1%) was added to quench the digestion. Finally, peptides were desalted by Sep-Pak *C*_18_ prior to Fe-IMAC enrichment.

The individual buffer conditions for the different samples are as follows. (1) BSA (1 mg/mL in 1xPBS, pH 7) was incubated with 1 mM of PhoX. (2) Protein Mixture Standard, consisting of alcohol dehydrogenase (baker’s yeast), myoglobin (equine heart), cytochrome C (equine heart), catalase (bovine), L-glutamic dehydrogenase (bovine liver) (each 1 mg/mL in 1xPBS, pH 7), was incubated with 1 mM of PhoX. (3) A HeLa cell pellet (5e7 of cells) was resuspended in ice-cold lysis buffer (700 µL, 50 mM HEPES, 150 mM NaCl, 1.5 mM MgCl_2_, 0.5 mM DTT, 1% benzonase, cOmplete Mini Protease inhibitor tablet) and soft lysis was performed by 30 to 40 quick pushes through a 27¾-gauge syringe. Then, cell debris was removed through centrifugation at 13,800 g for 10 min at 4 °C. The supernatant was incubated with 1 mM of PhoX for 1 h at r.t. Then, urea was added at a concentration of 8 M, followed by incubation with DTT (final concentration of 2 mM) for 30 min at 37 °C and alkylation with IAA (final concentration of 4 mM) for 30 min at 37 °C. This reaction was quenched by addition of DTT (final concentration of 2 mM). Then the sample was diluted four times with AmBic (50 mM, pH 8.3) and digested by incubation with LysC (1:75 enzyme to protein) and Trypsin (1:50 enzyme to protein) for 10 h at 37 °C, after which formic acid (1%) was added to quench the digestion. Finally, peptides were desalted by Sep-Pak *C*_18_ prior to both Fe-IMAC enrichment as well as LC-MS analysis. Phosphatase treatment of HeLa cell lysate peptides was applied as follows. Desalted peptides were dissolved at a concentration of 3 µg/µL in 1x CutSmart buffer (NEB, 50 mM potassium acetate, 20 mM tris-acetate, 10 mM magnesium acetate, 100 µg/mL BSA, pH 7.9). A volume of 2.4 µL of Alkaline phosphatase, calf intestinal (CIP, NEB, 10000 units/mL) was added and the mixture incubated at 37 °C overnight with shaking. Peptides were desalted using Sep-Pak *C*_18_.

Cross-linked peptides were enriched with Fe(III)-NTA cartridges primed at a flow rate of 100 µL/min with 250 µL of priming buffer (0.1% TFA, 99.9% ACN) and equilibrated at a flow-rate of 50 µL/min with 250 µL of loading buffer (0.1% TFA, 80% ACN). The flowthrough was collected into a separate plate. Dried samples were dissolved in 200 µL of loading buffer and loaded at a flow rate of 5 µL/min onto the cartridge. Columns were washed with 250 µL of loading buffer at a flow-rate of 20 µL/min and cross-linked peptides were eluted with 35 µL of 10% ammonia directly into 35 µL of 10% formic acid. Samples were dried down and stored at 4 °C until further use. Prior to LC–MS/MS analysis, the samples were resuspended in 10% formic acid.

### Data acquisition

Peptides were either directly infused through a nanospray emitter or were separated by nanoUHPLC (nanoElute, Bruker) on a 25 cm, 75 µm ID *C*_18_ column with integrated nanospray emitter (Odyssey/Aurora, ionopticks, Melbourne) at a flow rate of 250 nl/min. LC mobile phases A and B were water with 0.1% formic acid (v/v) and ACN with formic acid 0.1% (v/v), respectively. Samples were loaded directly on the analytical column at a constant pressure of 800 bar. In 70 min experiments, the gradient was kept at 0% B for 1 min, increased to 2% B over the next minute, followed by an increase from 2% to 34% B over 68 minutes. For column wash, solvent B concentration was increased to 85% for a further 8 minutes and kept at that concentration for an additional 12 minutes followed by re-equilibration to buffer A. For experiments at different gradient lengths, the time between 2% and 34% B was modified accordingly.

Data acquisition on the timsTOF Pro was performed using otofControl 6.0. Starting from the PASEF method optimized for standard proteomics, the following parameters were adapted: allowed charge states for PASEF precursors were restricted to 2-8. The base values for mobility dependent collision energy ramping were set to 85 eV at an inverse reduced mobility (1/K0) of 1.63 Vs/cm^2^ and 25 eV at 0.73 Vs/cm^2^; collision energies were linearly interpolated between these two 1/K0 values and kept constant above or below these base points (see Results and Discussion for more details). Each PASEF MSMS frame consisted of two merged TIMS scans acquired at 85% and 115% of the collision energy profile. To increase spectral quality, we set the target intensity per individual PASEF precursor to 40,000. For filtering PASEF precursors based on collisional cross section (CCS) and monoisotopic mass instead of 1/K0 and *m/z*, a modified acquisition implementation was used that transformed all potential precursor into CCS versus monoisotopic mass and applied a userdefined polygon as filter. We distinguish between “PASEF” where this filter is turned off (*i.e.* the standard acquisition approach) and caps-PASEF where this filter is turned on.

### Data analysis

The fragmentation spectra from all precursors with charge-state ≥ 2 were extracted from the recorded Bruker. d format files and stored in Mascot Generic Format (MGF) files with an in-house developed tool. The conversion procedure consists of two steps. (1) In the first step, fragmentation spectra of the same precursor are combined into a single spectrum. Matching of the precursors is performed with the following tolerances: precursor *m/z* +/− 20 ppm, retention time +/− 45 sec, and mobility +/− 2.5%. For each individual spectrum the noise level is estimated as the maximum intensity of the lowest 5% of peaks in the spectrum (for spectra with fewer than 10 peaks the noise level is fixed at 0.1). Combination of the spectra is achieved by clustering all peaks over all spectra within +/− 20 ppm of each other into a single peak with intensity equaling the sum of the combined individual peaks and the average signal-to-noise level of all spectra. (2) In the second step, each combined spectrum is de-isotoped (isotopes are reduced to a single peak at *m/z* of charge state of 1)(17), filtered for signal-to-noise of at least 1.5, and TopX filtered at 20 peaks per 100 Th. Together with the conversion procedure, an MGF-meta file is automatically created containing information on the Precursor Intensity, Mobility (1/K0), CCS, and monoisotopic Mass. The CCS values are calculated according to the Mason-Schamp equation(18); parameters are set to: temperature of 305 K, and the molecular weight of *N*_2_.

The MGF files for the synthetic peptides were annotated with in-house tooling using the same functionality as XlinkX version 2.4.0.193 to in-silico generate fragment peaks. The MGF files for the remaining experiments were analyzed with the same version(19). All database searches were performed against a FASTA containing the proteins under investigation supplemented with a contaminants list of 200 commonly detected proteins. For linear peptides, a database search was performed using Mascot version 2.7.0.0(20). Cysteine carbamidomethylation was set as fixed modification. Methionine oxidation and protein N-terminal acetylation were set as dynamic modifications. For the search of potential mono-linked peptides, water-quenched (*C*_8_*H*_5_*O*_6_*P*) and Tris-quenched (*C*_12_*H*_14_*O*_8_*PN*) were set as dynamic modifications. Trypsin was specified as the cleavage enzyme with a minimum peptide length of six and up to two missed cleavages were allowed. Filtering at 1% false discovery rate (FDR) at the peptide level was applied through Percolator(21). For cross-linked peptides, a database search was performed with PhoX (*C*_8_*H*_3_*O*_5_*P*) set as the cross-link modification. Cysteine carbamidomethylation was set as a fixed modification and methionine oxidation and protein N-terminal acetylation were set as dynamic modifications. Trypsin was specified as digestion enzyme and up to two missed cleavages were allowed. Furthermore, identifications were only accepted with a minimal score of 40 and a minimal delta score of 4. Otherwise, standard settings were applied. Filtering at 1% FDR at the peptide level was applied through a target/decoy strategy. Upon final assembly of the data, the protein identifications are FDR controlled to 1% and the identified cross-linked peptides are finally grouped on protein position. Further downstream analysis and visual representation of the results was performed with the R scripting and statistical environment(22) using ggplot(23) for data visualization.

### Data availability

All raw-files and the result files are described in Supplementary Table S1. The mass spectrometry proteomics data have been deposited to the ProteomeXchange Consortium via the PRIDE partner repository(24) with the dataset identifier PXD018189 (Username; Password: ikk1Pulf).

## Results and discussion

### Collisional energy optimization

As cross-linked peptides are different from unmodified peptides previously optimized settings potentially do not apply and we attempted to specifically optimize the fragmentation conditions for the identification of cross-linked peptides(25, 26). To determine the optimal collision energies for fragmenting cross-linked peptide pairs on the timsTOF Pro, we directly infused cross-linked synthetic peptides, isolated and independently subjected the ions at both charge state 2 (1/K0 = 0.98 Vs/cm^2^) as well as charge state 3 (1/K0 = 1.40 Vs/cm^2^) to fragmentation energies ranging from 10 – 100 eV in steps of 10 eV. After annotating the spectra, we optimized on the number of fragments as well as the production of a cross-link specific immonium ion from the lysine of peptide *α* connected via the cross-linker to the unfragmented peptide*β* (25). The optimal energy was determined from this analysis for the doubly charged cross-linked peptide at 70 eV and for the triply charged cross-linked peptide at 40 eV (see Supplementary Fig. S1A and B). We scaled the collision energy based on the collected 1/K0 values and according to the curve provided in Supplementary Fig. S1C. This curve limits the minimum collision energy to 20 eV, as below this range typically no fragmentation is observed for linear peptides, and the maximum collision energy to 80 eV, as at this energy typically over-fragmentation starts to be observed for linear peptides.

To investigate whether this initial curve is suitable for fragmentation of all cross-linked peptides, we additionally ran our BSA standard in multiple runs where each run used a fixed collision energy. The energies range from 20 to 120 eV in steps of 10 eV. From this collection of runs, a set of 496 identifications was covered by all fragmentation energies. After extraction of the sequence coverage for each spectrum, resulting in a sequence coverage trace, the traces of all peptide-pairs were correlated against all other peptide-pairs for each charge state independently. From the resulting heatmaps clusters could be defined, where peptide-pairs with the same behavior are grouped (Supplementary Fig. S2A and B). We found the main factor for distinguishing the clusters was mobility. The extracted optima fit reasonably well with the previously determined behavioral curve except for charge state 2 (Supplementary Fig. S2C). As the Bruker control software (otofControl) currently does not support breaking the calibration curve into different charge states, we opted to keep the parameters as previously determined, although we note that performance could possibly be improved with more advanced real-time acquisition logic.

### Mono-linked and cross-linked peptides can be distinguished by their behavior in ion mobility

Next, we cross-linked BSA and enriched for PhoX-linked mono- and cross-linked peptides by IMAC. From the BSA run recorded with PASEF, 192 linear peptides (of which 123 are mono-linked peptides; the remaining 69 are unspecifically binding to the IMAC material) and 80 cross-linked peptide-pair spectra were identified from a total of 17,516 separate scans (*i.e.* most spectra remain unidentified owing to the low precursor intensity thresholds employed in PASEF mode). When visualizing the mobility of the identified ions versus *m/z* (excluding unidentified), it is clear a degree of physical separation between the two classes of peptides is present (Fig. 1C). To gain insight into the resolution of this separation, a linear support vector machine (SVM) model was optimized to maximize the separation between mono-linked and cross-linked peptides; the distances of each identification to the linear model were then calculated (see Supplementary Fig. S3A-C). The density plot of the calculated distances indeed demonstrates there is a clear physical separation between the two classes of ions, showing that the extra dimension of ion mobility assists in improving the level of detection (Fig. 1D).

On top of the physical separation, mono-linked peptides can potentially also be excluded from sequencing. However, a large degree of overlap with the cross-linked peptides hampers the differentiation between mono-linked and cross-linked peptides by the data acquisition software. Translation to CCS values and visualization against *m/z* demonstrates that the charge state 2 mono-link identifications separate from cross-link identifications (Fig. 1E). Moreover, lower *m/z* regions of the higher charge states were uniquely identified as mono-link and separate from the higher *m/z* regions of the higher charge states that were identified as cross-links. However, it is not yet trivial to make the separation between the classes of molecules and therefore we sought for a way to improve this further.

### CCS assisted precursor selection improves PASEF for crosslinked peptides

First, to visually show the separation, we further translated the *m/z* values depicted on the x-axis (Fig. 2A) to monoisotopic mass (Fig. 2B). Here the mono-linked peptides cluster in the bottom-left corner while the cross-linked peptides cluster in the top-right corner. The separation between these two classes hinges on a CCS of 500 *Å*^2^ and a monoisotopic mass of 2 kDa, above which a polygonal area (Supplementary Table S2) can be drawn that encapsulates most of the cross-linked peptides while excluding most of the mono-linked peptides (Fig. 2B; red dotted polygon). Counting the precursors selected for fragmentation (Fig. 2C) resulted in 7784 unidentified fragmentation spectra outside and 9460 unidentified fragmentation spectra inside the polygon suggesting that these are normally distributed and for the vast majority genuine noise. For the mono-linked peptide identifications, only 19 out of 192 fall inside the polygon, representing a 91 % reduction in these identifications if those outside of the polygon were to be excluded. In sharp contrast, we only detect a single identification outside the polygon for the cross-linked peptides.

**Figure 2.**
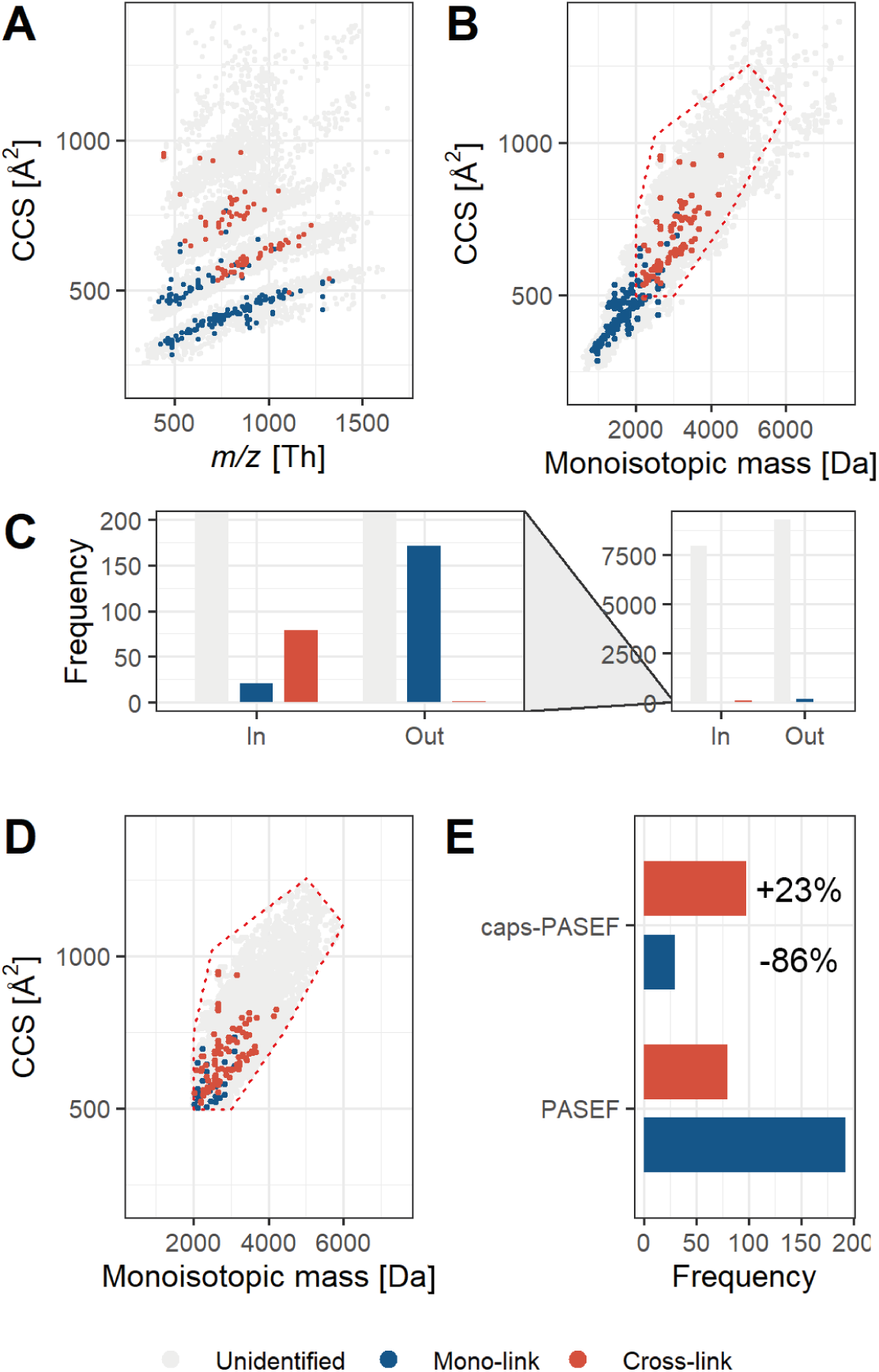
Collisional Cross Section Assisted Precursor Selection (caps-PASEF) applied to cross-linked BSA. **(A)** Identical to Fig. 1A; distribution of *m/z* (Th) versus CCS for all classes of ions, converted to **(B)** monoisotopic mass versus CCS. The red dotted polygon contains most of the cross-linked peptides.**(C)** Overview of the classes of ions in- and outside the defined polygon area. **(D)** Data collected in optimized caps-PASEF mode (‘Unidentified’: likely noise; ‘Mono-link’: 30; ‘Cross-link’: 98). **(E)** Comparison of mono-link and cross-link identification results in PASEF and caps-PASEF mode. The ratio cross-linked versus mono-linked peptides increases from 0.4:1 to 3.2:1 going from PASEF to caps-PASEF mode.

Based on these results, we hypothesized that the observed separation can provide the basis for an effective data acquisition protocol, whereby the mass spectrometer is focused to predominantly sequence the cross-linked peptides – a strategy we termed collisional cross section assisted precursor selection PASEF or caps-PASEF. Detection of isotope patterns and the consequent translation to monoisotopic mass as well as CCS can efficiently be performed in real-time by the data acquisition software; a higher degree of errors is anticipated for calling the 12*C* peak, but this will not have a large impact on these values and we expect them to be sufficiently precise. This protocol was integrated into the on-line data acquisition software (otofControl) and programmed with the polygon displayed in Figure 2B. From the BSA run recorded in caps-PASEF mode it is clear the mass spectrometer is solely sequencing precursors within the programmed polygon (as visualized in Fig. 2D). From the caps-PASEF run, 30 peptides (of which 23 mono-linked) and 98 cross-linked peptides are detected. This represents an almost 85% reduction in identified mono-linked peptides when comparing against the PASEF run. Excitingly, a substantial increase of ~20% in the number of cross-linked peptides is observed. We cannot exclude from this data, obtained for a single protein, that this is within statistical variation, although the increase is somewhat supported by the modest increase of 21 to 30 mono-link identifications within the polygon.

### Application of caps-PASEF to medium complexity samples

To verify whether caps-PASEF works well with the sample complexities typically analyzed by XL-MS (purified protein complexes of three or more subunits), we analyzed a standard protein mixture of six proteins with PASEF and caps-PASEF, applying in both cases the abovementioned optimized parameters (Fig. 3). Inspection of the physical separation as performed for the BSA dataset shows that even though the sample complexity increased, the TIMS device is still able to physically separate the mono-linked from the crosslinked peptides (Fig. 3A). The distribution of normal, cross-linked and unidentified peptides is similar to the one observed in the BSA dataset (Fig. 3B). From the bar charts at the top and on the right, it can clearly be observed that, as before, the mono-linked peptides are shifted to the bottom/left compared to cross-linked peptides in the overall distributions; eliminating the bottom/left regions will enable the mass spectrometer to predominantly sequence cross-linked peptides. Applying caps-PASEF with the previously defined polygon for the cross-linked BSA samples successfully prevents sequencing of the peptide background (Fig. 3C). By copying the polygon from the BSA run, a few low molecular weight cross-linked peptides are excluded as well. As these however constitute cross-linked peptides of short sequence lengths, these identifications tend to be problematic for high complexity mixtures and elimination can potentially assist to reduce false positive rates(27).

**Figure 3.**
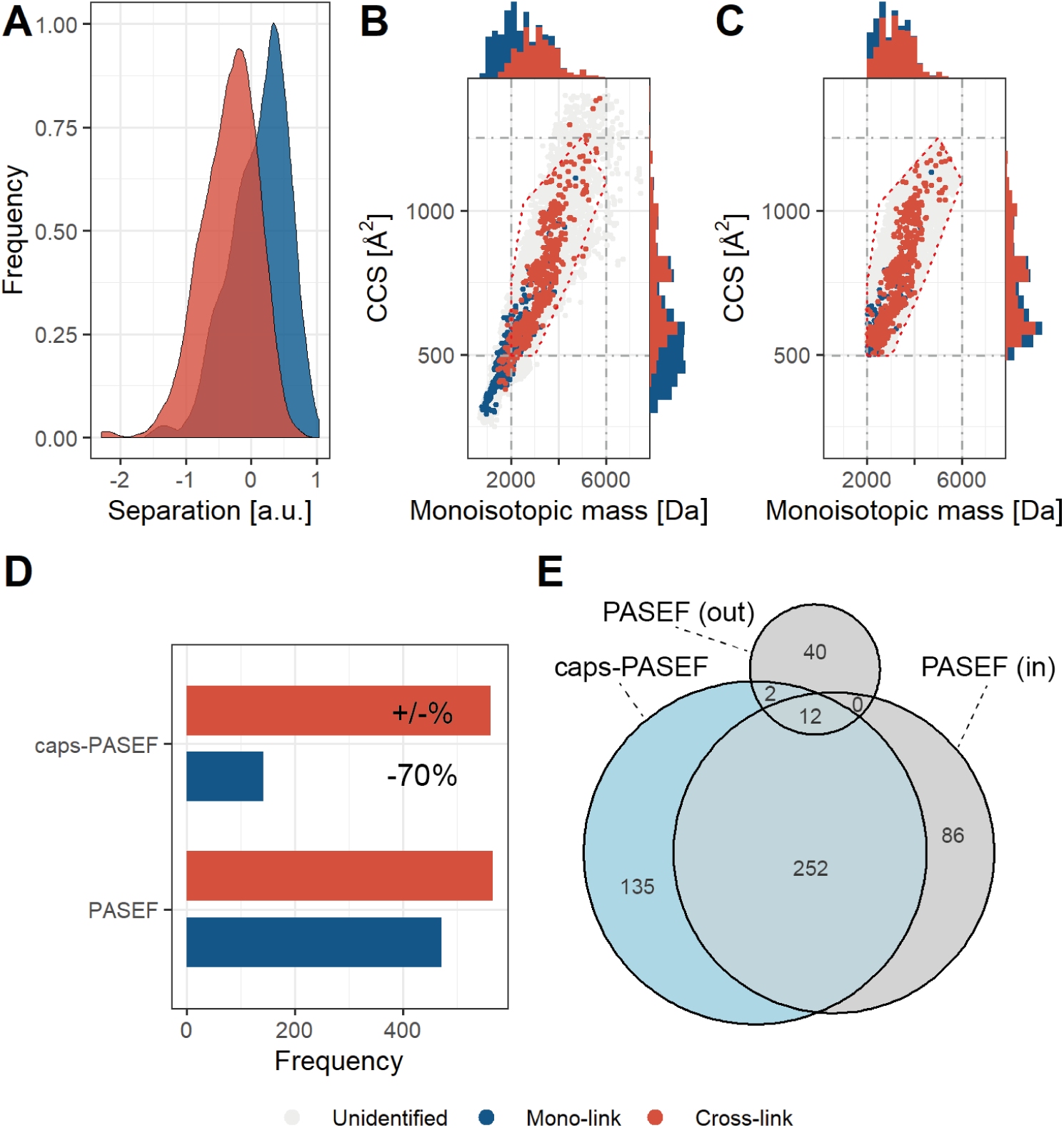
Collisional Cross Section Assisted Precursor Selection (caps-PASEF) applied on a standard protein mixture. **(A)** Physical separation in mobility space, in arbitrary units, for cross-linked versus mono-linked peptides. **(B)** Diagram displaying the monoisotopic mass versus CCS in PASEF mode (‘Unidentified’: likely noise; ‘Mono-link’: 472; ‘Cross-link’: 566). **(C)** Diagram displaying the monoisotopic mass versus CCS in caps-PASEF mode (‘Unidentified’: likely noise; ‘Mono-link’: 143; ‘Cross-link’: 562). **(D)** Comparison of mono-link and cross-link identification results in PASEF and caps-PASEF mode. The ratio cross-linked versus mono-linked peptides increases from 1.2:1 to 3.9:1 going from PASEF to caps-PASEF mode. **(E)** Overlap in detected cross-linked peptides between PASEF (’in’ denotes inside and ‘out’ denotes outside the polygon) and caps-PASEF.

Comparing the identification results of the two different runs shows that caps-PASEF identifies close to the same number of crosslinked peptides as the PASEF run (PASEF: 566; caps-PASEF: 562), while reducing the amount of mono-link identifications by ~70% (PASEF: 472; caps-PASEF: 143) (Fig. 3D). Inspection of the sequences of the identifications shows an overlap of ~75% between the measurements when considering the identifications within the defined polygon (Fig. 3E). A subset of 14 identifications were found in the PASEF run outside the polygon (of which 12 were also identified inside the polygon), which can be attributed to variation in the CCS values derived from the TIMS device which can fluctuate in most cases by a maximum of 10% potentially driving the identification outside the polygon (Supplementary Fig. S4A). Effects incurred by variations in the mass detection are not anticipated (Supplementary Fig. S4B). Interestingly, caps-PASEF identifies 49 additional crosslinked peptides inside the polygon; an increase of ~15% revealing that by focusing the acquisition to a region of interest more data of interest can be acquired.

### Application of caps-PASEF to proteome-wide cross-linking

Application of PASEF to a PhoX enriched cross-linked full cellular lysate shows that physical separation of the different classes of formed peptides is progressively more difficult and will likely not bring additional depth in identifications if the complexity becomes too high (Fig. 4A). To verify whether our caps-PASEF approach still brings benefit at this level of complexity, we inspected the distribution of the identifications of normal, cross-linked and unidentified peptides, and found it similar to those observed in the BSA and the protein mix datasets, although much more overlap occurs between the mono-link and cross-link identifications (Fig. 4B). Application of caps-PASEF with the same polygon as used before indeed successfully removes a large majority of the mono-link identifications (Fig. 4C). Comparing the identification results of the two different runs shows that caps-PASEF identifies ~10% more cross-linked peptides as the PASEF run (PASEF: 332; caps-PASEF: 364), while reducing the amount of mono-link identifications by ~60% (PASEF: 3606; caps-PASEF: 1581) (Fig. 4D). Inspection of the sequences of the identifications shows an overlap of ~60% between the measurements when considering the identifications within the defined polygon (Fig. 4E).

**Figure 4.**
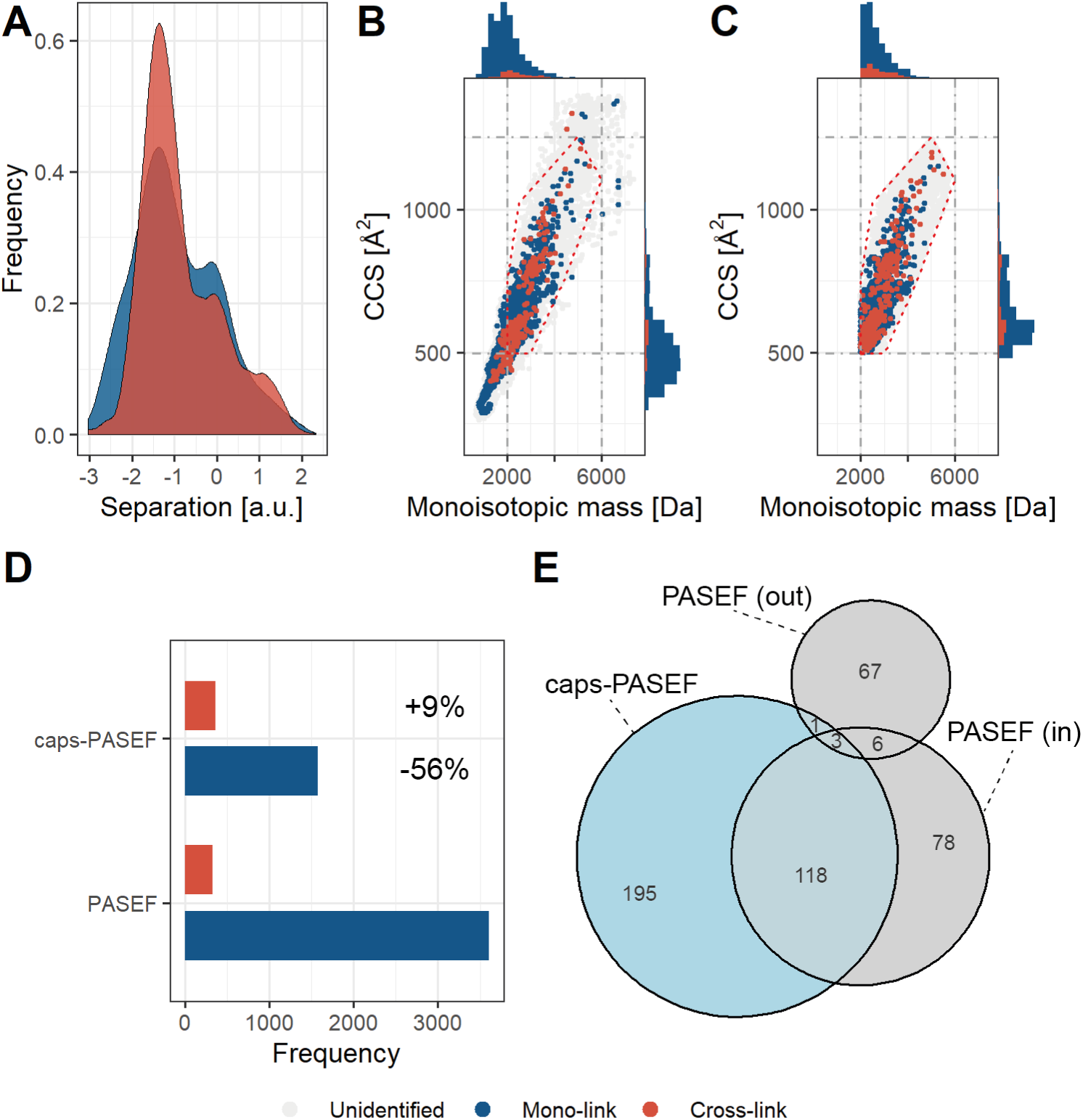
Collisional Cross Section Assisted Precursor Selection (caps-PASEF) applied to a complex cross-linked Hela cell lysate following PhoX enrichment. **(A)** Physical separation in mobility space, in arbitrary units, for cross-linked versus mono-linked peptides. (B) Diagram displaying the monoisotopic mass versus CCS in PASEF mode (‘Unidentified’: likely noise; ‘Mono-link’: 3606; ‘Cross-link’: 332). **(C)** Diagram displaying the monoisotopic mass versus CCS in caps-PASEF mode (‘Unidentified’: likely noise; ‘Mono-link’: 1581; ‘Cross-link’: 364). **(C)** Comparison of mono-link and cross-link identification results in PASEF and caps-PASEF mode. **(D)** The ratio cross-linked versus mono-linked peptides increases from 0.1:1 to 0.2:1 going from PASEF to caps-PASEF mode. **(E)** Overlap in detected cross-linked peptides between PASEF (‘in’ denotes inside and ‘out’ denotes outside the polygon) and caps-PASEF.

Overall, even though the benefit for proteome-wide cross-linking is reduced to approximately 10%, the focusing of the mass spectrometer on the ions of interest will have large benefits. Further improvements to the mass spectrometry platforms will result in higher quality fragmentation scans and therefore better identifications. For all experiments, the mono-link background was heavily reduced while not affecting the cross-link identifications and in some cases markedly improving the number of cross-link identifications. This effect was even more pronounced when observing the number of identifications within the polygon, for which we observed improvements of 20 – 50 %.

## Conclusions

XL-MS represents a powerful approach to uncover structural details of proteins and protein-complexes(10, 11), even in highly complex samples(9). Despite its power, the technique has however suffered from limited analytical depth due to the low reaction efficiency of the used reagents. With the introduction of enrichable cross-linking reagents like PhoX(9, 28), this can partly be resolved. With these reagents the sample complexity can be reduced, focusing only on peptides modified by the cross-linking reagent, forming mono-linked and cross-linked peptide products. Further improvements are however still required to fully unlock the potential of XL-MS, as the mono-linked peptides do not provide the sought after structural information and typically make up more than half of the sample load after enrichment. Here, we described the development of a novel acquisition approach utilizing ion mobility to physically separate the mono-linked from the cross-linked peptides, providing better signal-to-noise to the latter class of ions. Additionally, we present a novel acquisition technique capable of preventing sequencing of a large majority of mono-linked peptides, while still sequencing the desired cross-linked peptides. The approach is exemplified on the Bruker timsTOF Pro, which incorporates a trapped ion mobility device in a mass spectrometry platform geared towards shotgun proteomics. From the acquired data we have demonstrated that the data acquisition software can make the required a-priori distinction between mono-linked and cross-linked peptides. This focusses the acquisition, a feature largely beneficial for complex mixtures.

With the availability of enrichable cross-linkers and data acquisition protocols as described here, we envision that XL-MS can outgrow the extraction of structural information from highly purified samples. Even though the experimental conditions have been developed, the latter will still require statistical approaches to interpret the detected cross-linked peptides and what they truly represent. Notwithstanding, we believe the future for XL-MS is particularly bright.

## Acknowledgments

We thank all members of the Hecklab for their helpful discussions and contributions. We acknowledge support from the Netherlands Organization for Scientific Research (NWO) funding the Netherlands Proteomics Centre through the X-omics Road Map program (project 184.034.019). This work is supported through the European Union Horizon 2020 program INFRAIA project Epic-XS (Project 823839), and a seed grant kindly provided by the Utrecht Institute for Pharmaceutical Sciences (UIPS). All authors critically read and edited the manuscript.

## Contributions

R.A.S. and A.J.R.H. conceived of the study. B.S. performed the cross-linking experiments and performed together with R.A.S. the data analysis. H.W.P.T. and R.A.S. developed the raw data extraction and performed data analysis. J.F.G. assisted with the interpretation of the ion mobility data. E.B. analyzed the fragmentation energy data. M.L. and B.S. acquired the data. R.A.S. and O.R. conceived of the caps-PASEF strategy.

## Notes

O.R. and M.L. are employees of Bruker Daltonik GmbH, manufacturer of the timsTOF Pro, and thus disclose competing financial interests.

