## Supplemental Material for "Benefits of Collisional Cross Section Assisted Precursor Selection (caps-PASEF) for Cross-linking Mass Spectrometry"

**Running title:** The timsTOF Pro applied to XL-MS

**Supplementary Table S1 – RAW files used in this study.** File-type ‘mgfmeta’ contains CCS, mobility and intensity. This is linked to file-type ‘mgf’ on scannumber/first scan. Files ending with _PSMS.txt contain the Mascot results and files ending with _CSMS.txt contain the XlinkX results.

| | Associated files | Type | Panels | Comments |
|---|---|---|---|---|
| **Fig.** | BSAPhoX60minlike60_Slot1-1_01_3657.d | PASEF | **1B-E,** | This set of files |
| **1,2,** | BSAPhoX60minlike60_Slot1-1_01_3657.mgf | | **S3A-C** | contains |
| **S3,** | BSAPhoX60minlike60_Slot1-1_01_3657.mgfmeta | | **S4A,B** | acquisition on |
| **S4** | BSAPhoX60minlike60_Slot1-1_01_3657_PSMs.txt | | | BSA from which |
| | BSAPhoX60minlike60_Slot1-1_01_3657_CSMs.txt | | | the polygon for |
| | | | | the further runs is |
| | BSAPhoX60minlike60_polygon_Slot1-1_01_3669.d | caps-PASEF | **1B-E,** | based. |
| | BSAPhoX60minlike60_polygon_Slot1-1_01_3669.mgfmeta | | **2A-E,** | |
| | BSAPhoX60minlike60_polygon_Slot1-1_01_3669_PSMS.txt | | **S4A,B** | |
| | BSAPhoX60minlike60_polygon_Slot1-1_01_3669_CSMS.txt | | | |
| **Fig.** | ProteinMixPhoX70minlike60_Slot1-3_01_3671.d | PASEF | **3A-D** | This set of files |
| **3** | ProteinMixPhoX70minlike60_Slot1-3_01_3671.mgf | | | contains the |
| | ProteinMixPhoX70minlike60_Slot1-3_01_3671.mgfmeta | | | acquisition on our |
| | ProteinMixPhoX70minlike60_Slot1-3_01_3671_CSMs.txt | | | protein mix. |
| | ProteinMixPhoX70minlike60_Slot1-3_01_3671_PSMs.txt | | | |
| | ProteinMixPhoX70minlike60_polygon_Slot1-3_01_3672.d | caps-PASEF | | |
| | ProteinMixPhoX70minlike60_polygon_Slot1-3_01_3672.mgf | | | |
| | ProteinMixPhoX70minlike60_polygon_Slot1-3_01_3672.mgfmeta | | | |
| | ProteinMixPhoX70minlike60_polygon_Slot1-3_01_3672_CSMs.txt | | | |
| | ProteinMixPhoX70minlike60_polygon_Slot1-3_01_3672_PSMs.txt | | | |
| **Fig.** | HeLaPhoX150minlike60_16MSMS_Slot1-4_01_3676.d | PASEF | **4A-D** | This set of files |
| **4** | HeLaPhoX150minlike60_16MSMS_Slot1-4_01_3676.mgf | | | contains the |
| | HeLaPhoX150minlike60_16MSMS_Slot1-4_01_3676.mgfmeta | | | acquisition on |
| | HeLaPhoX150minlike60_16MSMS_Slot1-4_01_3676_PSMs.txt | | | HeLa. |
| | HeLaPhoX150minlike60_16MSMS_Slot1-4_01_3676_CSMs.txt | | | |
| | HeLaPhoX150minlike60_polygon16MSMS0_75_Slot1-4_01_3677.d | caps-PASEF | | |
| | HeLaPhoX150minlike60_polygon16MSMS0_75_Slot1-4_01_3677.mgf | | | |
| | HeLaPhoX150minlike60_polygon16MSMS0_75_Slot1-4_01_3677.mgfmeta | | | |
| | HeLaPhoX150minlike60_polygon16MSMS0_75_Slot1-4_01_3677_PSMs.txt | | | |
| | HeLaPhoX150minlike60_polygon16MSMS0_75_Slot1-4_01_3677_CSMs.txt | | | |
| **Fig.** | 20190520_AAAAKAAAAAR_726_10-90eV_d3-110_PASEF.d | PASEF direct | **S1A-C** | Initial definition of |
| **S1** | 20190520_AAAAKAAAAAR_726_10-90eV_d3-110_PASEF_CSMs.txt | infusion | | the CE calibration |
| | | | | curves |
| | 20190520_AAAAKAAAAAR_1089_10-90eV_d3-110_PASEF.d | | | |
| | 20190520_AAAAKAAAAAR_1089_10-90eV_d3-110_PASEF_CSMs.txt | | | |
| **Fig.** | BSAPhoX60min20_Slot1-2_01_3659.d | PASEF fixed | **S2** | This set of files |
| **S2** | BSAPhoX60min20_Slot1-2_01_3659.mgf | collision | | contains the |
| | BSAPhoX60min30_Slot1-2_01_3661.d | energies | | acquisitions on |
| | BSAPhoX60min30_Slot1-2_01_3661.mgf | ranging from | | BSA with fixed |
| | BSAPhoX60min40_Slot1-2_01_3663.d | 20 to 120 in | | collision energies |
| | BSAPhoX60min40_Slot1-2_01_3663.mgf | steps of 10 | | acquired in |
| | BSAPhoX60min50_Slot1-2_01_3665.d | | | scrambled order. |
| | BSAPhoX60min50_Slot1-2_01_3665.mgf | | | |
| | BSAPhoX60min60_Slot1-2_01_3667.d | | | The CCS values |
| | BSAPhoX60min60_Slot1-2_01_3667.mgf | | | were read directly |
| | BSAPhoX60min70_Slot1-2_01_3668.d | | | from the MGF file |
| | BSAPhoX60min70_Slot1-2_01_3668.mgf | | | (in the header: |
| | BSAPhoX60min80_Slot1-2_01_3666.d | | | TITLE). The file |
| | BSAPhoX60min80_Slot1-2_01_3666.mgf | | | csmtable.txt |
| | BSAPhoX60min90_Slot1-2_01_3664.d | | | contains the |
| | BSAPhoX60min90_Slot1-2_01_3664.mgf | | | identifications |
| | BSAPhoX60min100_Slot1-2_01_3662.d | | | linked to the |
| | BSAPhoX60min100_Slot1-2_01_3662.mgf | | | scannumbers and |
| | BSAPhoX60min110_Slot1-2_01_3660.d | | | csmtable.mgf all |
| | BSAPhoX60min110_Slot1-2_01_3660.mgf | | | spectra extracted |
| | BSAPhoX60min120_Slot1-2_01_3658.d | | | from all mgf file. |
| | BSAPhoX60min120_Slot1-2_01_3658.mgf | | | |
| | csmtable.txt | | | |
| | csmtable.mgf | | | |

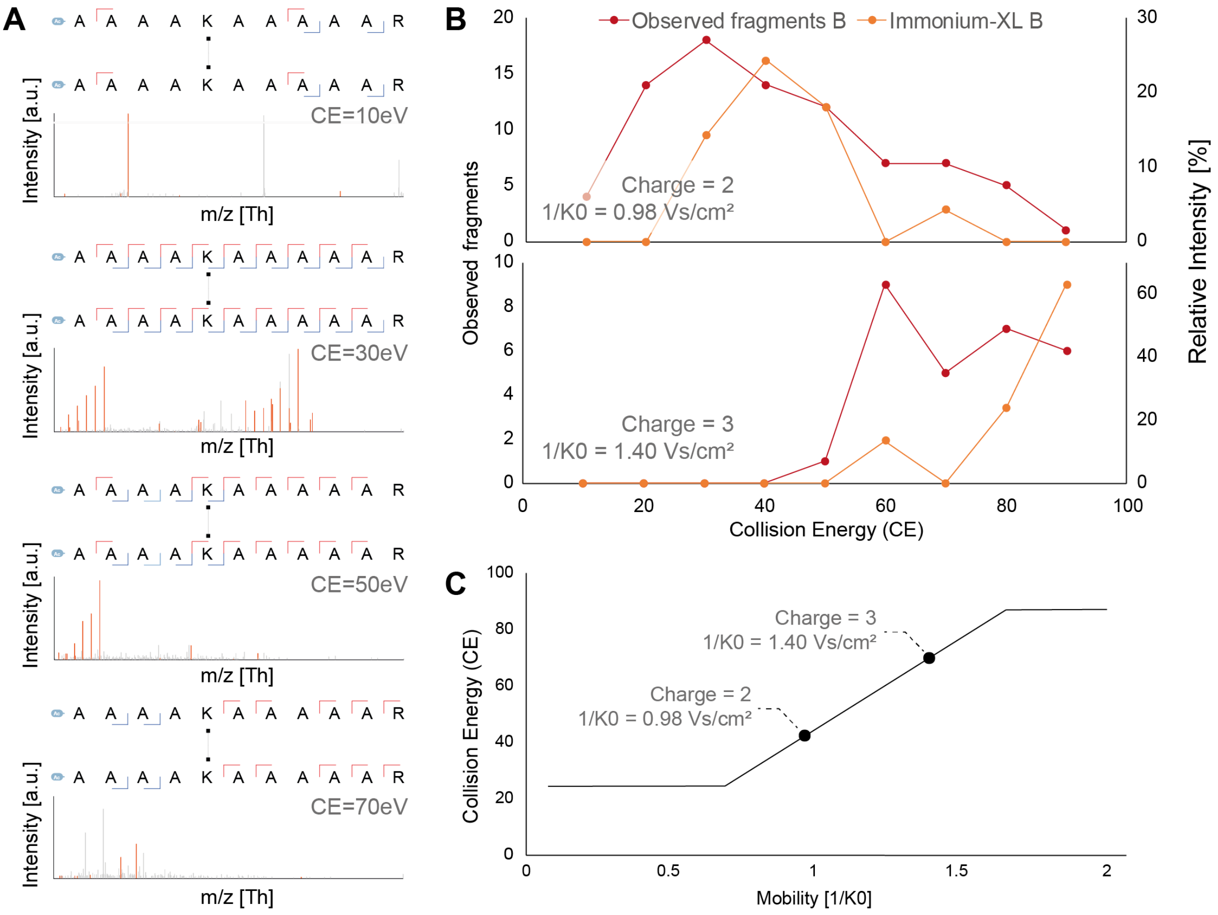


**Supplementary Figure S1 – Collision energy optimization on the cross-linked peptide dimer Ac-AAAAKAAAAAR. (A)** Fragmentation spectra for the cross-linked peptide dimer at charge state 2 at different collision energies. At low collision energies, no fragments are produced, which only start to appear at energies that are more elevated. At energy levels higher than the optimal, over-fragmentation starts to occur, visible here in the disappearance of fragments linking an intact peptide to the other, fragmented, peptide. **(B)** Expressing the quality of the fragmentation spectra at various collision energies in observed fragments (equates directly to sequence coverage) and relative intensity of cross-linking reagent specific immonium ion (see Steigenberger *et al*(1)) the optimal points for charge state 2 is found at ~ 40 eV and for charge state 3 at ~70 eV. **(C)** Final calibration curve linearly extrapolating based on the two observations and limiting the energy at 25 eV, below which typically no fragmentation is observed, and at 85 eV, above which typically only over fragmentation is observed.

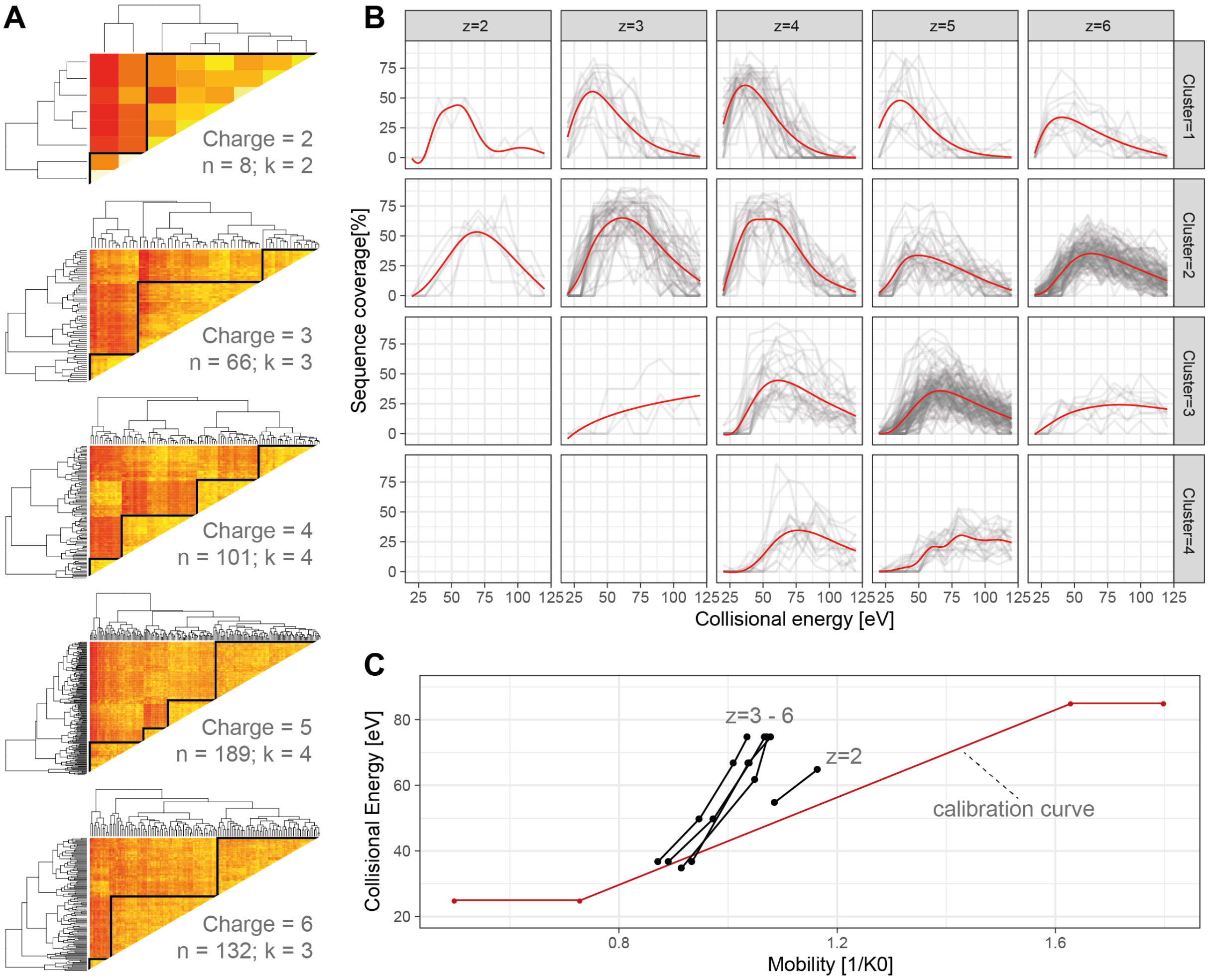


**Supplementary Figure S2 – Collision energy optimization on BSA with fixed collision energies. (A)** After extraction of identifiable spectra (no FDR control was applied, spectra for collisional energies where the identification could not be made were located by correlating the spectra with precursor mass within 20 ppm and retention time within 2 min), for all spectra the sequence coverage was calculated – resulting in a sequence coverage trace over all the collisional energies. By correlating each trace against all other traces for each charge state independently, we were able to construct dendrograms from which matching traces could be selected in an unbiased fashion. **(B)** Extraction of the clusters shows that the unsupervised clustering method extracts sequence coverage traces with similar properties and that the apex of the average trace (indicated by the red line) slowly shifts to higher collisional energies for each cluster. **(C)** Plotting the apex collisional energy for the median mobility of each cluster shows that the initially determined calibration curve for lower mobilities fits well for charge state 3 - 6, although for higher mobilities starts to diverge from the calibration. For charge 2 we observer divergent behaviour from the rest of the charge states, suggesting that altering the calibration curves in the instrument control software could potentially have a beneficial effect on the identification performance. As the control software does not at the point of writing not support the complex behaviour as extracted by this analysis, we opted to keep the initially extracted calibration curve as a conservative estimate that fits well with the lowest charge state.

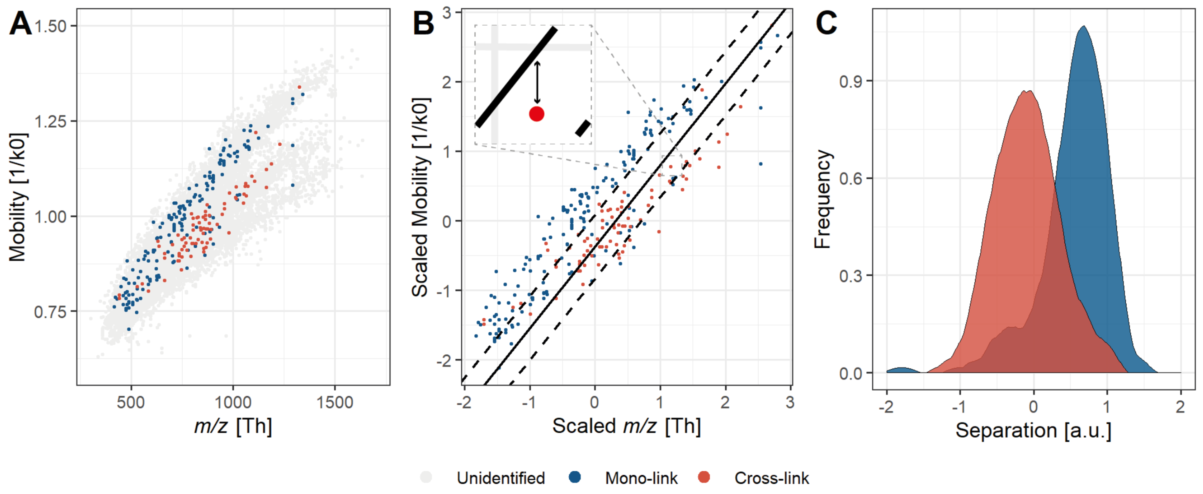


**Supplementary Figure S3 – Estimation of physical separation in mobility space for the BSA run in PASEF mode. (A)** The m/z (Th) versus Mobility (1/K0) for all classes of molecules. **(B)** After removal of the unidentified molecules and z-score scaling of the values a linear support vector (SVM) was fit, optimized on separating between the peptides/mono-links and the cross-links, in R with the package 'e1071'. The solid line denotes the separating plane fit by the SVM and the dotted lines the confidence interval. **(C)** The mobility distance (*i.e.* on the y-axis alone) of each identification to the solid line is calculated and expressed in the density plot for both peptide/mono-link and cross-link identifications.

**Supplementary Table S2 – The polygon definition for our caps-PASEF experiments.**

| Mono-isotopic mass | CCS |
|---|---|
| 2000 | 500 |
| 2000 | 750 |
| 2500 | 1000 |
| 5000 | 1200 |
| 6000 | 1100 |
| 4400 | 750 |
| 3000 | 500 |

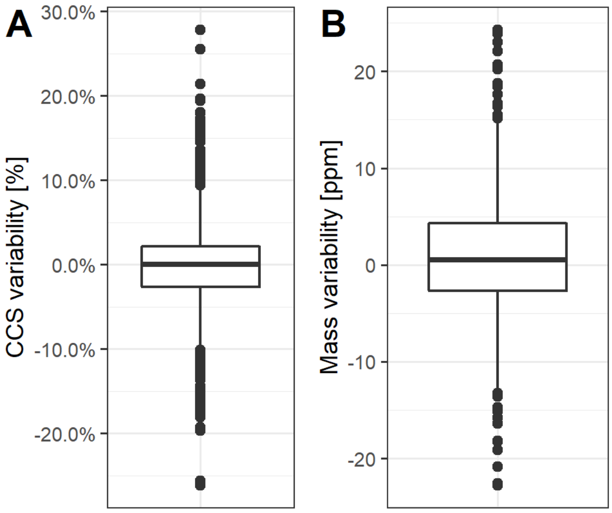


**Supplementary Figure S4 – Inter-measurement variability for the protein-mix. (A)** Detection of CCS values for identified cross-linked peptides shows the TIMS device has a precision of +/- 10% over two measurements for the vast majority of the cases, with 50% of the detections within +/- 2.5%. **(B)** Deviations between the detected mono-isotopic masses of the two measurements confirms that the instrument measures with an accuracy of +/- 20 PPM.
